# Auditory Speech Processing Efficiency: Functional Connectivity Graph Theory Analysis of Lexical Processing Effort

**DOI:** 10.64898/2025.12.04.690545

**Authors:** Paulina Skolasinska, Adam T. Eggebrecht, Julia L. Evans

**Affiliations:** McGill University, Montreal, QC, CA; Washington University in St. Louis, St. Louis MO, USA; University of Texas at Dallas, Dallas, TX, USA

**Keywords:** spoken word processing, functional near-infrared spectroscopy, functional connectivity, graph theory, mental lexicon, linguistic efficiency

## Abstract

**Purpose:** Network science models of real-time spoken word processing predict that word frequency and sublexical phonotactic probability directly influence verbal working memory. Behavioral accuracy and speed of processing suggest that high frequency words differ in cognitive processing effort as compared to low frequency words. Cognitive processing effort, assessed indirectly through changes in prefrontal oxygenated (HbO) and deoxygenated (HbR) hemoglobin concentrations, also suggest that high and low frequency words differ in cognitive effort. This novel study examines potential differences in linguistic processing effort and efficiency directly, using measures of functional connectivity (FC) strength, modularity, and global efficiency, derived from the listener’s prefrontal cortex hemodynamic responses using a paradigmatic verbal working memory paradigm.

**Method:** A total of 20 neurologically typical participants (age 18-21 years) completed an auditory *n*-back working memory task comparing performance for words differing in whole-word frequency and sub lexical phonotactic probability. Changes in HbO and HbR hemoglobin concentration were recorded with a continuous-wave, multi-channel fNIRS system (TechEn, Inc., Milford, MA) using a 20-channel optode montage across the prefrontal cortex. Partial correlation coefficients were calculated between each channel pair to produce 20 x 20 FC matrices for each participant. Prefrontal networks were constructed as a graph where nodes in the graph are the fNIRS measurements and edges are partial correlations between pairs of measurements.

**Results:** Word frequency effects were evident in both in the behavioral and functional connectivity measures. Task performance, regardless of word frequency, was related to brain measures, such that higher prefrontal FC strength was negatively correlated with accuracy (*d’*) and high modularity was negatively correlated with response times for low frequency words. In keeping with the direction of the brain-behavior correlations, the highest performing participant had lower FC strength and higher efficiency whereas the lowest performing participant had higher modularity and lower global efficiency.

**Conclusions:** We identified network properties that are linked to lexical indices of cognitive processing effort in typical participants. These preliminary results point to strategies that could assess language function in atypical populations in future studies.

## Introduction

Emergentist psycholinguistic models assume that real-time spoken sentence comprehension occurs at the intersection of different syntactic representations and cognitive mechanisms such as verbal working memory capacity, attentional control, updating, and inhibition (Montogomery et al., 2009). This framework views comprehension as a process whereby the listener must rapidly build phrase and clause structures across the incoming words to construct a syntactic–semantic representation of sentence meaning. Gibson and colleagues (Gibson, et al., 2019) argue that if the primary purpose of language is the transferring of information in the context of social interactions, then the structure of languages should have evolved to prioritize efficiency of use. From an information theory framework, this means that the form of all languages should have evolved so that the ideal speaker can transmit and the ideal listener can receive information successfully with minimal effort.

This predicts not only that linguistic efficiency should be measurable across all linguistic levels (e.g., lexical, syntactic, etc.) but that efficiency and linguistic cognitive processing effort should be linked. From this perspective a linguistic structure’s utility can be examined with respect to the ease with which a speaker can encode a message into the speech signal and the ease with which the listener can subsequently decode the message. At the level of the lexicon, behavioral measures such as reaction time (e.g., speed) and accuracy (e.g., d-prime, *d’*) have traditionally been viewed as indirect measures of processing effort.

Network science models of real-time lexical processing assume that as the listener processes the incoming speech signal, the initial sound of a word activates a set of all possible word candidates which compete until enough of the incoming speech stream allows the listener to narrow in on the actual word (e.g., TRACE, cohort, and neighborhood activation models; Luce & Pisoni, 1998; Marslen-Wilson & Tyler, 1980; Marslen-Wilson & Zwitserlood, 1989; McClelland & Elman, 1986). The underling structure of the mental lexicon directly influences this spreading activation process (e.g., Vitevitch, et al, 2012; Vitevitch, 2019) and the ease and efficiency efficiency of bottom-up activation spreading activation from the acoustic input (Law & Edwards, 2015) and lateral inhibition connections, which enable words that have strong activations to directly suppress less active competitors, make word processing more or less effortful and efficient (Gur & McMurray, 2025).

These lexical network models use nodes to represent words in the mental lexicon, and edges to connect nodes (e.g., words) that are phonologically related (Vitevitch, 2008). They show that words having a low degree of phonological overlap with other words (e.g., degree of phonological neighborhood density, Vitevitch & Luce, 2016) influence listeners speed and accuracy in spoken word recognition; where words having low phonological overlap are processed faster and more accurately by listeners than words having a high phonological overlap with other words in the lexicon (Chan & Vitevitch, 2009).

The overall frequency with which a word occurs in a language (e,g., word frequency) as well as the frequency with which different phoneme combinations appear in words within a given language (e.g., phonotactic frequency/probability) also influence the speed and accuracy with which listeners can hold words in verbal working memory (e.g., Gathercole, Frankish, Pickering, & Peaker, 1999; Roodenrys, Hulme, Lethbridge, Hinton, & Ninmo, 2002; Storkel & Rogers, 2000; Van Overschelde, 2002). Word frequency, followed by phonotactic frequency also predicts listeners’ word recall from short-term memory (e.g., Evans, et al., 2022; Luce & Pisoni, 1998; Luce & Large, 2001; Mainela-Arnold & Evans, 2005; Mainela-Arnold, Evans & Coady, 2010; Vitevitch and Luce, 1999).

Neural correlates of processing efficiency derived from functional MRI (fMRI), EEG, and functional near-infrared spectroscopy (fNIRS) have suggest differences in processing effort on word frequency and phonotactic probability (e.g., Evans, et al., 2022; Weiss, et al., 2023). For example, EEG lexical decision studies, indirectly suggest that processing effort is lower for high-word frequency/phonotactic probability (HF) as comparted to low-word frequency/phonotactic probability (LF) spoken words (Evans, et al., 2022), with mean N400 amplitude beginning approximately 500 ms after the word onset and lasting through the end of the epoch for low as compared to high frequency words showed a greater negative defection along with a second pattern of greater negative defection of LF and compared to HF words during the 700-1200 ms time window. Researchers have also used functional near infrared spectroscopy (fNIRS) and an *n*-back working memory design to examine more directly the cognitive effort inherent in activation, updating, and inhibiting words differing in word frequency and phonotactic probability (Berglund-Barraza, et al., 2019, 2020).

In our prior work (Berglund-Barraza, et al., 2019, 2020) we used fNIRS and a novel n-back working memory design to examine the cognitive effort inherent in activation, updating, and inhibiting words differing in word frequency and phonotactic probability. Using Chan & Vitevitch’s (2009) lexical network theoretical framework, we hypothesized that the combined influence of word frequency and phonotactic probability would have an even greater effect on cognitive processing effort due to the combined effect of having both higher resting activation levels and lower access thresholds, which would translate directly into greater processing efficiency and lower cognitive effort. Specifically, we predicted that this ease of processing would be evidenced by higher behavioral accuracy, faster speed of processing, and that overall HbO and HbR concentration in the frontal/prefrontal cortex being lower for the high word + high phonotactic probability (HWF+PP) words as compared to the low word + low phonotactic probability (LWF+PP) words (which we refer to as high frequency = HF and low frequency = LF for simplicity). By leveraging the experimental control provided by the *n*-back task we were able to measure the effect of word frequency and phonotactic probability the storage demands required by the listener to actively maintain and update the working memory. Consistent with our predictions, listeners were faster and more accurate in processing/updating/ actively maintaining HF words as compared to LF words. Participants’ accuracy for HF words was also significantly correlated with the degree of HbO changes in the lateral inferior region of the right hemisphere, and reaction time was correlated with HbO in the right lateral inferior and medial frontal regions.

A limitation of the Berglund-Barraza et al. study design is that we used overall concentrations of HbO and HbR hemoglobin as a proxy measure of cognitive effort. More nuanced measures derived from functional connectivity (FC) and graph theory might be better suited to answer questions on language processing efficiency. FC represents the temporal correlation between hemodynamic or electrical activity in different brain regions during a task or at rest, indicating that those regions are functionally linked and may work together to perform specific cognitive, perceptual, or motor tasks (Biswal et al., 1995). Hemodynamic FC can be measured using various neuroimaging techniques including fMRI and fNIRS (Oldham & Fornito, 2018). Graph theory provides a powerful framework for quantifying different properties of such networks (Fornito et al., 2016).

Like many naturally occurring networked systems, the human brain is organized in a small-world and modular fashion, due to economical constraints driving minimization of connection cost (Achard & Bullmore, 2007; Bullmore & Sporns, 2012). Such an organization scheme ensures efficient information transfer in the entire network through a balance of specialized local processing and long connections facilitating efficient global integration (Rubinov & Sporns, 2011; Bertolero et al., 2015). Specifically, a graph theoretical measure of modularity captures the strength of segregation between non-overlapping functional modules. Optimal modularity depends on the cognitive context, where dominance of local, segregated processing is favored at rest and during easy cognitive tasks, while higher integration due to a need for inter-module information exchange is required as cognitive task demands grow. For example, in an fMRI study of typical young adults, Cohen and D’Esposito (2016) showed lower modularity (or greater network integration) during the *n*-back task compared to a motor sequence tapping task and resting-state. Such greater network integration was correlated with high accuracy on the *n*-back task.

At the same time, network integration is metabolically costly and is therefore avoided when possible. Measures capturing global integration such as average network FC, might then inform about maximizing processing efficiency while minimizing energetic cost. Inefficient FC patterns have been previously described in multiple neurological disorders, including neurodegeneration (Chen et al., 2021; Yao et al., 2010), schizophrenia (Lynall et al., 2010), and autism spectrum disorder (ASD; Rudie et al., 2012). Graph-theoretical studies using MRI have identified low connectivity within brain networks stemming from weaker short-range and long-range connections, as well strong connectivity between the networks, resulting in low modularity and high global efficiency at the whole-brain scale in ASD compared to typically developing peers. In contrast, other developmental disorders, like the developmental language disorder (DLD) has not been extensively studied using graph theoretical approaches.

However, even in typical development and young adulthood, individual differences in cognitive performance are related to network properties that may represent processing efficiency. For example, structural brain networks, which provide the underlying architecture for their functional counterpart (Honey et al., 2009), were shown to become more segregated and efficient in typical development due to strong connections of the central (or hub) nodes. Crucially, increased modularity mediated improvements in executive performance with age (in a sample of 8-22 years old; Baum et al., 2017), highlighting the importance of the network-based approaches in the study of complex cognition.

### Purpose

In this study, we analyzed the data from our prior fNIRS studies (Berglund-Barraza et al., 2019; 2020) focusing on graph theory measures of FC strength and network modularity in a group of young adults having typical language ability. This enabled us to compare the cognitive processing effort and efficiency required to process, update, and actively maintain words having high word frequency and high phonotactic probability in verbal working memory as compared to words having low word frequency and low phonotactic probability. Additionally, we characterized the spatial distribution of network hubs that were engaged during the task to determine if FC strength and modularity were sensitive to individual differences in working memory performance across the participants.

## Materials and Methods

### Experimental Design

In our previous studies (e.g., Berglund-Barraza et al., 2019; 2020) we leveraged the experimental control of the *n*-back task to examine potential differences in spoken word processing effort for words differing in whole word frequency and phonotactic probability. In our previous study we held the working memory load constant by having the participants complete a *2*-back auditory working memory task where they were asked to determine if the target item is the same as the item that was 2-back in the sequence. By using a 2-back task paradigm this allowed us to keep the demands of the task constant while manipulating the demands of the spoken words themselves while manipulating the demands on the participants to actively maintain each spoken word in verbal working memory while simultaneously update the incoming information and inhibiting lexical competitors after each trial to keep track of the order of the last *n* items in the sequence (Owen et al., 2005; Jaeggi et al., 2010).

Participants completed a total of 14 blocks (7 High, 7 Low) where they heard a series of one-syllable real words differing in whole word frequency and phonotactic probability (**Figure 1**). Because word frequency and phonotactic probability differentially influence the ability of the listener to inhibit lexical cohort interference effects (Coady et al., 2010; Mainela-Arnold, et al., 2008; Mainela-Arnold, et al., 2010) we compared word frequency + phonotactic probability at the same time across the seven blocks where the words were high word frequency + high phonotactic probability (the “HF” blocks) and seven blocks where the words were words low word frequency + low phonotactic probability (the “LF” blocks). Using a conventional fMRI block design, the seven HF and seven LF blocks were presented in an alternating manner beginning with a HF block.

**Figure 1.**
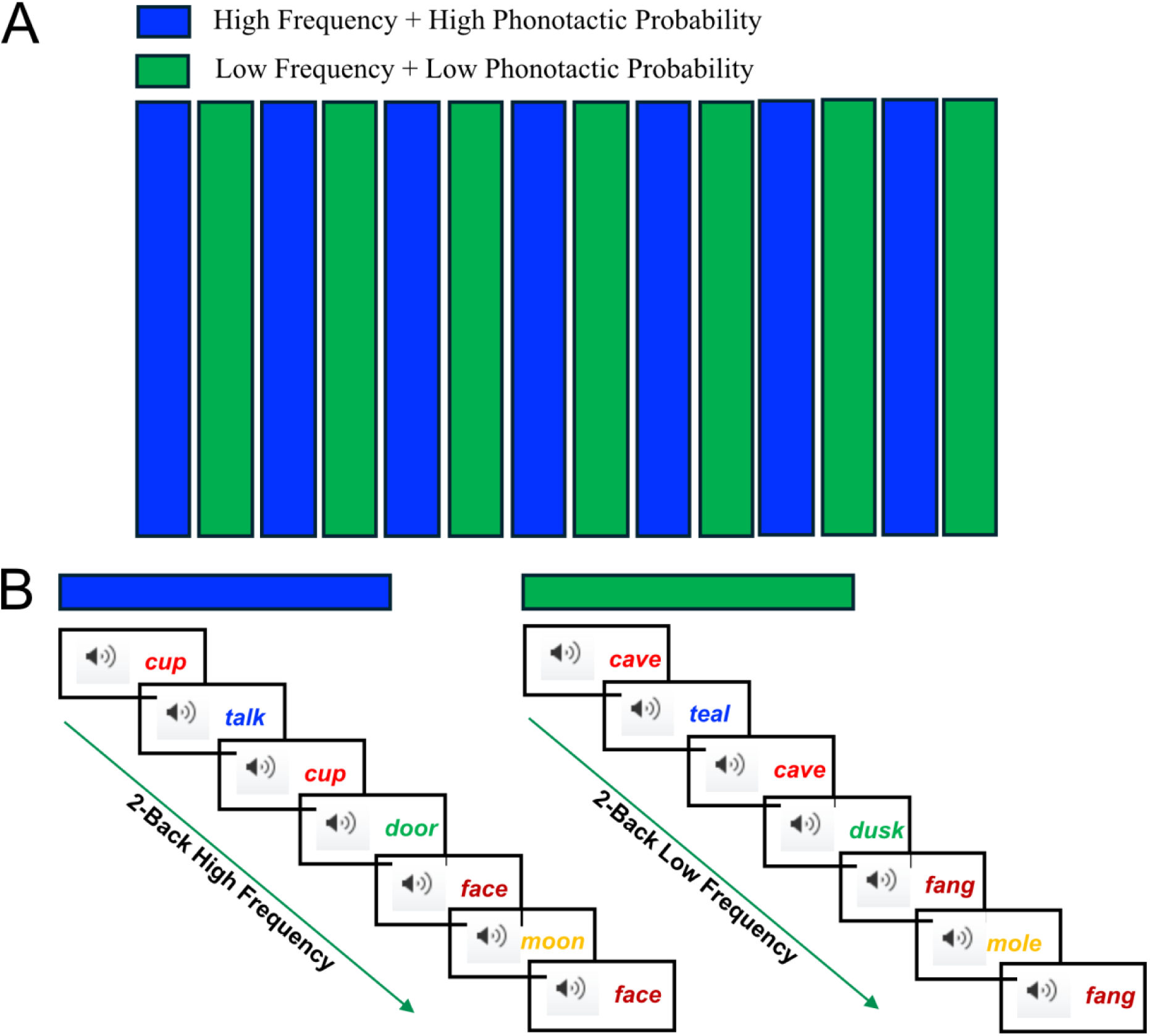
Design of the 2-back auditory working memory task. Block order (A) and stimulus presentation (B). A. High Frequency + High Phonotactic Probability B. Low Frequency + Low Phonotactic Probability

We created seven different fixed random-order sequences of the hits and correct rejections that were assigned to each of the seven HF and LF blocks so the serial order of hits/correct rejections did not differ across those blocks. The hit rate was the same for all the blocks (25%) and no more than three consecutive hits occurred in a row. All blocks contained a total of 98 words. The initial phoneme of target words within each of the blocks was controlled so that the initial phoneme of the congruent target/non-target pairs differed. The lists were also controlled so that the distribution of initial consonants in the blocks did not differ for the HF and LF lists. The word order in each block also constrained to ensure that there was no semantic priming effect between target/non-target words.

#### Stimuli

The stimuli consisted of one-syllable spoken real words digitally recorded at 44.1 Hz by an adult female native speaker of English. Each word was approximately 700-milliseconds in duration, and each sound file contained an initial 50 milliseconds of silence, and each word began with a 5-millisecond envelope. Silence was added at the end of each sound file so that all files were 1000-milliseconds in duration and words were presented with an interstimulus interval (ISI) of 500 milliseconds resulting in each block being 90 seconds in duration. Each block was separated by a 15 second resting state period of silence where the participants were instructed to rest quietly and not move.

The words in the HF and LF blocks differed in both whole word frequency and sublexical phonotactic frequency (word frequency, *F* (1, 13) = 310.5, *p* < .001, partial h^2^ = .353; biphone phonotactic frequency, *F* (1, 13) = 9877, *p* < .001). Because the inhibitory effect neighborhood density can the effect of sublexical phonotactic frequency, in part because the two measures are highly correlated (i.e., Coady, Evans, Kluender, 2010), all the words in both conditions had low neighborhood density with ratings at or below 20 and did not differ for the HF and LF blocks, *F* (1, 12) = 0.01 and *p* = .92. Imageability was also controlled, with all of the words in both conditions having imageability ratings at or above 5.0 (Walker & Hulme, 1999) and not differing for HF and LF blocks, *F* (1, 13) = 0.29 and *p* = .60. (See **Table 1**).

**Table 1.** Word frequency, sublexical phonotactic probability, imageability, and neighborhood density for the high frequency (HF) and low frequency (LF) blocks.

| Condition | Block | Word<br>Frequency<sup>a</sup> | Phonotactic<br>Probability<sup>b</sup> | Imageability<sup>c</sup> | Neighborhood<br>Density<sup>d</sup> |
| --- | --- | --- | --- | --- | --- |
| HF | 1 | 191.72 | 3.17 | 5.11 | 22.03 |
|  | 2 | 182.65 | 3.12 | 5.30 | 21.51 |
|  | 3 | 203.43 | 3.15 | 5.20 | 21.94 |
|  | 4 | 176.68 | 3.14 | 5.25 | 21.00 |
|  | 5 | 251.06 | 3.21 | 5.27 | 21.20 |
|  | 6 | 215.89 | 3.14 | 5.31 | 20.75 |
|  | <u>7</u> | <u>253.74</u> | <u>3.17</u> | <u>5.26</u> | <u>20.60</u> |
|  | Mean | 210.74* | 3.16* | 5.24<sup>ns</sup> | 21.29<sup>ns</sup> |
|  | SD | 28.96 | 0.03 | 0.06 | 0.52 |
| LF | 1 | 3.09 | 1.36 | 5.44 | 20.23 |
|  | 2 | 2.43 | 1.26 | 5.13 | 21.67 |
|  | 3 | 2.68 | 1.32 | 5.22 | 21.38 |
|  | 4 | 2.47 | 1.27 | 5.17 | 20.66 |
|  | 5 | 2.29 | 1.26 | 5.13 | 21.92 |
|  | 6 | 2.20 | 1.25 | 5.22 | 21.09 |
|  | <u>7</u> | <u>2.26</u> | <u>1.27</u> | <u>5.21</u> | <u>22.32</u> |
|  | Mean | 2.49* | 1.28* | 5.22<sup>ns</sup> | 21.32<sup>ns</sup> |
|  | SD | 0.29 | 0.04 | 0.10 | 0.67 |

<sup>a</sup>*Coltheart (1981) MRC Psycholinguistic Database;*
*http://www.psy.uwa.edu.au/mrcdatabase/uwa_mrc.htm.*
<sup>b</sup>*Vitevitch and Luce, 2004; http://www.bncdnet.ku.edu/cgi-bin/DEEC/post_ppc.vi.*
<sup>c</sup>*Washington University in St. Louis; http://128.252.27.56/Neighborhood/Home.asp.*
<sup>d</sup>*Cortese and Fugett, 2004, http://www.psychonomic.org/ARCHIVE/.*

#### Participants

A total of 21 typical language, females, ages 18-25, participated in the Bergland-Barraza (2019, 2020) studies. One participant was excluded due to excessive movement artifacts. A total of 20 of the participant’s data were analyzed for this study. All participants were right-handed, monolingual English speakers. No participant had a history of neurological injury or disease, seizure disorder, or psychiatric diagnosis, and no participant had received any speech language or resource services in school. All participants completed written informed consent protocols in accordance with the guidelines of the University of Texas at Dallas (UTD) Institutional Review Board, which approved the protocol.

#### Functional Near Infrared Spectroscopy (fNIRS)

The fNIRS scans were collected at the Callier Center for Communications Disorders in Dallas, a Research Center at UTD, in The Child Language and Cognitive Processes Laboratory under author JLE. A multichannel continuous-wave TechEn CW6 system (TechEn Inc., Milford, Massachusettes), with a sampling frequency of 25 Hz was used to collect the data. The system uses near-infrared lasers at λ = 690 nm (HbR) and λ = 830 (HbO) nm as emitters, and avalanche photodiodes (APDs) as detectors. A fiber optic probe that contained 6 emitting optodes and 12 detecting optodes with emitter-detector distances of 3 cm was placed bilaterally and symmetrically on the participant’s forehead (**Figure 2**). The probe was secured in place with an elastic band during the experiment. The bottom of the probe array was placed just above the eyebrows.

**Figure 2.**
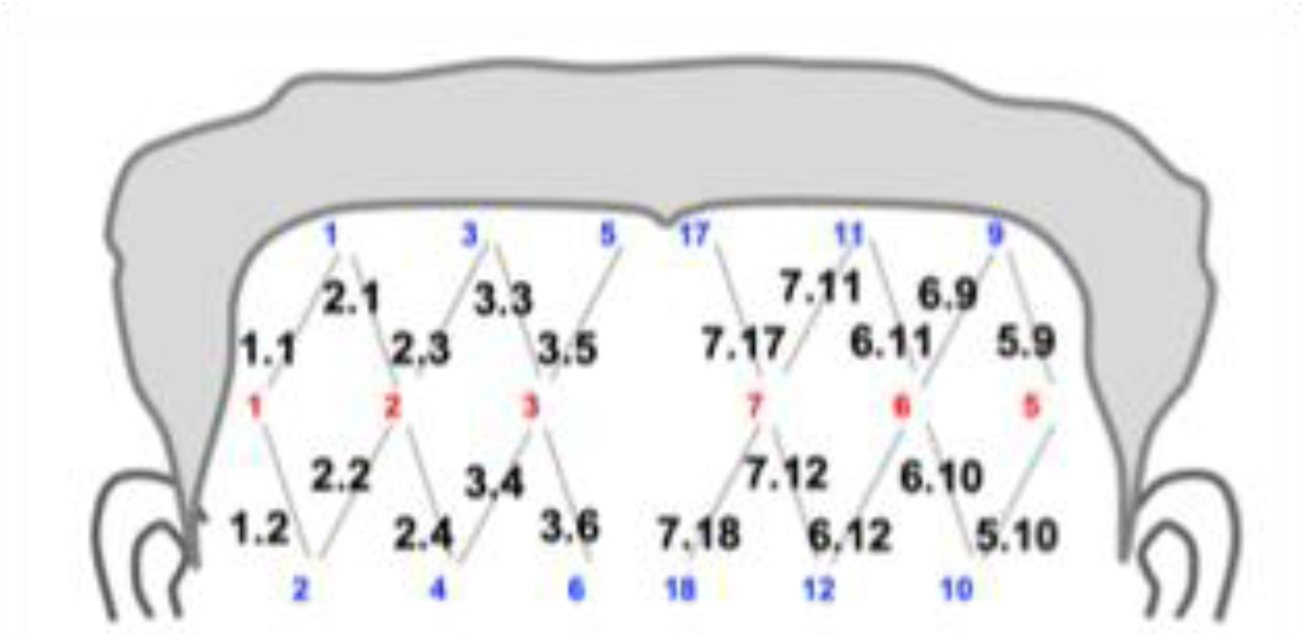
Optode layout. Emitters in red, detectors in blue.

#### Data Processing and Artifact Removal

Data were preprocessed using the NeuroDOT toolbox in MATLAB, including custom scripts. The raw nirs data and was logmean-transformed, detrended, and bandpass filtered between 0.008 – 0.09 Hz. The mean signal was regressed out from the data at each wavelength, followed by data resampling at 1 Hz. Next, partial correlation coefficients were calculated separately for HbO and HbR, as well as for HF and LF tasks, between each channel pair while controlling for the interactions between all other channel pairs (Marrelec et al., 2006). Such partial correlation matrices were Fisher r-to-z-transformed using the arctanh function.

The data were censored, removing timepoints with high motion, identified using the global variance of temporal derivatives (GVTD) metric (Sharafati et al., 2020). The GVTD threshold was set to 0.007. However, we excluded one participant who was missing 50%+ of the HF or LF data after censoring. Noisy channels were not removed to maintain the structure of the correlation matrices.

#### Graph-Theory Analysis

Following preprocessing, four 20 x 20 FC matrices were obtained for each participant, one for each signal type (HbO_2_, HbR) and word frequency (HF, LF). The average strength of prefrontal FC (“FC strength”) was calculated by averaging the correlation coefficients between all pairs of brain regions using unthresholded matrices. Network modularity (*Q*) was also calculated, using the Newman’s algorithm (Newman, 2006),

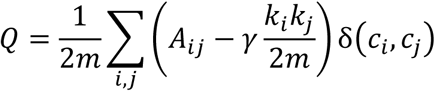

which is a weighted sum of each edge in the input matrix *A* with rows *i* and columns *j* minus the degree *k* of *i* and *j* scaled by the resolution parameter gamma (*γ* = 1) and the number of edges *m*. *c_i_* and *c_j_* give module assignments for each pair of nodes, so that the delta function equals 1 if *c_i_* = *c_j_* and 0 otherwise.

The optimal community structure of a graph was obtained by maximizing the modularity measure over all possible partitions, by maximizing within-group connections and minimizing between-group connections. Therefore, modularity reflected the strength of the separation of non-overlapping clusters of nodes (i.e., modules) within the entire network. Modularity was calculated using thresholded matrices (Rubinov & Sporns, 2010). As there is no single agreed upon threshold, we used multiple thresholds, that is from 1% to 99%, with the increment of 1%, with the % determining the proportion of edges to be retained, resulting in weighted, undirected networks having varying sparsity. The matrices obtained using thresholds of 10% and below were excluded due to excessive graph sparsity.

Global efficiency was also calculated over this threshold range (Latora & Marchiori, 2001; Vragović et al., 2005). Global efficiency is the average of the inverse shortest path distances between all node pairs,

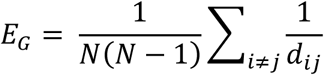

where *N* is the number of network nodes, here represented by channels, and *d* is the shortest path distance from node *i* to node *j.* Intuitively, global efficiency represents the general ease of information transfer over the entire network. Computation of graph theoretical metrics was performed using R package igraph (Csárdi & Nepusz, 2006).

#### Statistical Analysis

Our measure of performance accuracy was *d*′, a sensitivity index in *z* units, that differentiates the means for the signal (i.e., target) versus noise (i.e., foils) and is a direct measure of the listener’s ability to recognize/differentiate target from foil trial while controlling for listener sensitivity and response bias (MacMillan & Kaplan, 1985). The higher the *d’* value the better participant performance (i.e., accuracy in recognizing matches and avoiding foils). In this study perfect performance was *d*′ = 8.6. Reaction time (RT) was analyzed for correct trials with no trimming of outliers. Data were corrected using the Greenhouse-Geisser correction applied to the probability values to adjust for repeated measures. Repeated measures ANOVAs were conducted for *d*′ and RT as dependent measures with group as the between subject factor and frequency (HF, LF) as the within subject factor.

In addition, to examine individual differences across the participants, and the degree to which the highest and lowest performing participants’ ability to recognize/differentiate the target from the foil trials fell outside of the normal range, *z-*scores were calculated for *d*′, RT, FC strength, and modularity using the entire group’s performance as the normative database in the same manner as that described in Burgland-Barraza et al., (2020) and Brown et al., (2014). This allowed us to identify the participants who had the highest and the lowest average *d’,* respectively. The scores of the highest performing participant and the lowest performing participant were omitted when calculating the means and standard deviations used as reference in the calculation of the *z*-scores, so that each extreme score was not included to obtain the statistics that it was to be compared to 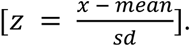. All measures were separated by frequency (HF, LF) while the brain metrics were additionally investigated by signal type (HbO, HbR). Modularity and efficiency measures were averaged over the range of thresholds (10% - 99%).

To test the effect of word frequency on FC measures, we used linear mixed models, while accounting for the threshold. The models included linear, and quadratic effects of the threshold, their interactions with word frequency, as well as individual intercept (*u_0i_*), individual slope (*u_1i_*) and individual random error (*ε_ij_*): *y_ij_ = β_0_ + β_1_word frequency_ij_ + β_2_threshold_ij_ + β_3_threshold ^2^ + β_5_(word frequency_ij_ ⋅ threshold_ij_) + β_6_(word frequency_ij_ ⋅ threshold_ij_^2^) + u_0i_ + u_1i_ threshold_ij_ + ε_ij_,* where *ij* corresponds to the *i*-th measurement of participant *j*. For these models, the threshold was considered as continuous. The *lmer* function from the R lme4 package was used for the linear mixed models, and the emmeans R package for used for the post-hoc comparisons with the Holm correction for multiple comparisons. Finally, we performed correlations between the brain and the behavioral measures. One-sided Pearson’s correlations tests were performed. We expected better performance (higher *d’* and lower RT) to be related to lower FC strength, indicative of higher efficiency, as well as lower modularity, and higher global efficiency, suggesting greater integration in support of cognitive performance in line with Cohen and D’Esposito (2016).

## Results

### Behavioral Measures

Accuracy as measured by *d’* over the course of the high- and low-frequency blocks are shown in **Figure 3A**. A repeated measures ANOVA revealed that the main effect of word-frequency approached significance, *F*(1,36) = 3.19, *p* = 0.08, partial η^2^ 0.08, power 0.41 where participants’ accuracy in correctly detecting the target word 2-back in the sequence was marginally better overall for the LF (*M* = 4.54) as compared to the HF (*M* = 3.82) blocks. For the main effect of block, *F*(6,216) = 3.15, *p* < 0.01, partial η^2^ 0.8, power .91, the quadratic effect was significant, *F*(1,36) = 10.49, *p* < 0.01, partial η^2^ 0.22, power 0.88, indicating that participants’ accuracy increased significantly until the middle of the task and then declined slightly toward the end of the experiment. The significant word frequency by block interaction effect, *F*(6,216) = 7.49, *p <* 0.01, partial η^2^ 0.17, power 1.0 indicates that over the course of the experiment, participants’ accuracy was greater for the LF as compared to the HF blocks.

**Figure 3.**
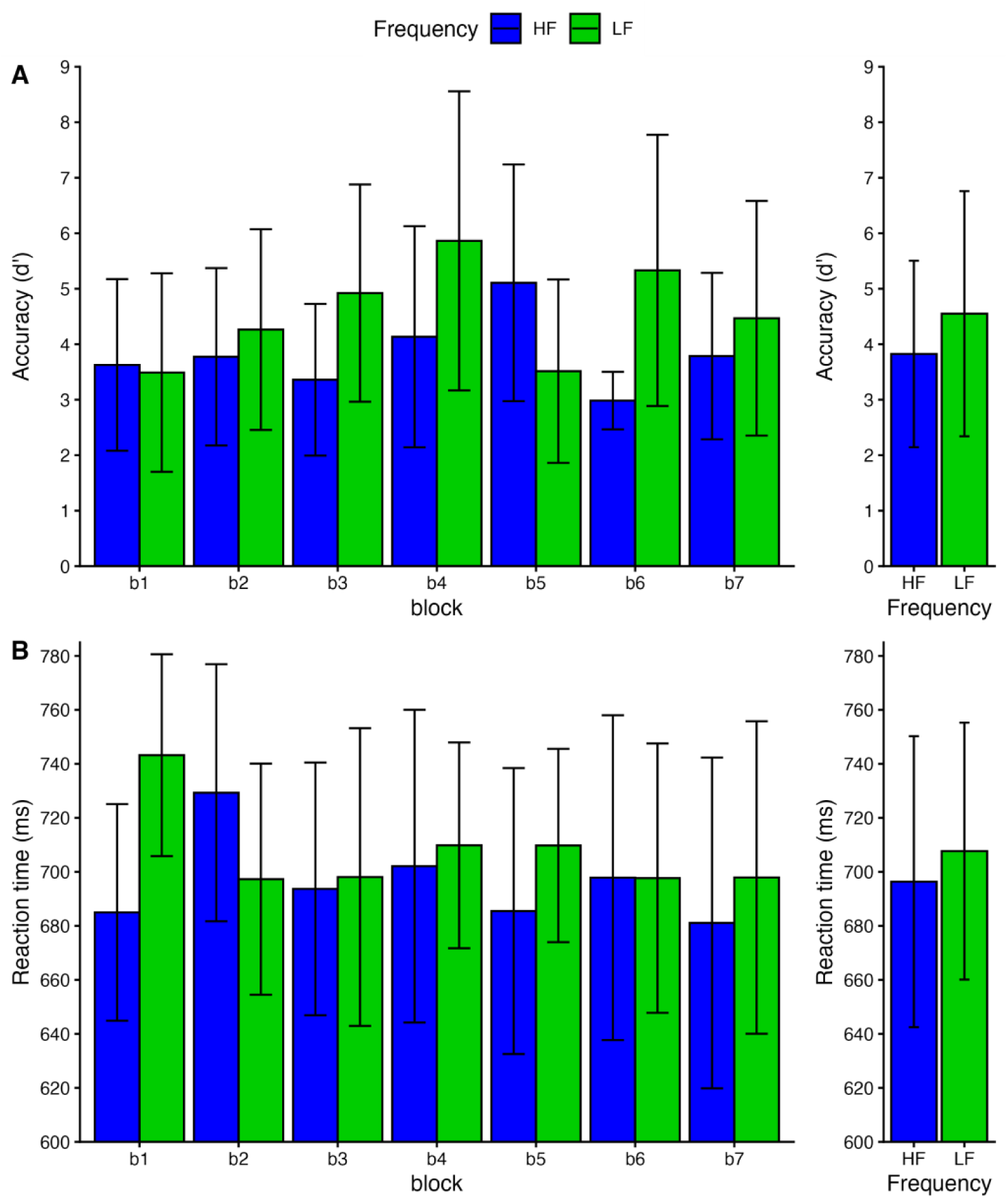
Accuracy in *d*-prime (A) and reaction time in milliseconds (RT; B) for high/low word frequency and phonotactic probability for each task block 1-7 (left) and averaged across the blocks (B). The error bars represent ± 1 standard deviation.

RTs for correct trials are shown in **Figure 3B**. A repeated measures ANOVA revealed that although the main effect of word frequency approached significance, *F*(1,36) = 1.012, *p =* 0.32, partial η^2^ 0.02, power 0.16, there was a significant frequency by block interaction *F*(3.7,133.7) = 4.83, *p =* 0.002, partial η^2^ 0.11, power 0.94 and a significant main effect of block, where the linear effect was significant *F*(1,36) = 6.33, *p =* 0.01, partial η^2^ 0.15 indicating that participants’ speed of response declined over the course of the experiment to a greater extent for the LF as compared to the HF blocks.

### Hemodynamic Network Measures

No main effect of word frequency was observed for FC strength [HbO: *t*(17) = -0.51, *p* = .62; HbR: *t*(17) = 1.00, *p* = .33], modularity [HbO: *t*(17) = -0.13, *p* = .90; HbR: [*t*(17) = -0.42, *p* = .68], global efficiency [HbO: *t*(17) = 0.42, *p* = .68; HbR: *t*(17) = -1.38, *p* = .19] when averaged across the thresholds. **See Figure 4** for boxplots of each measure.

**Figure 4.**
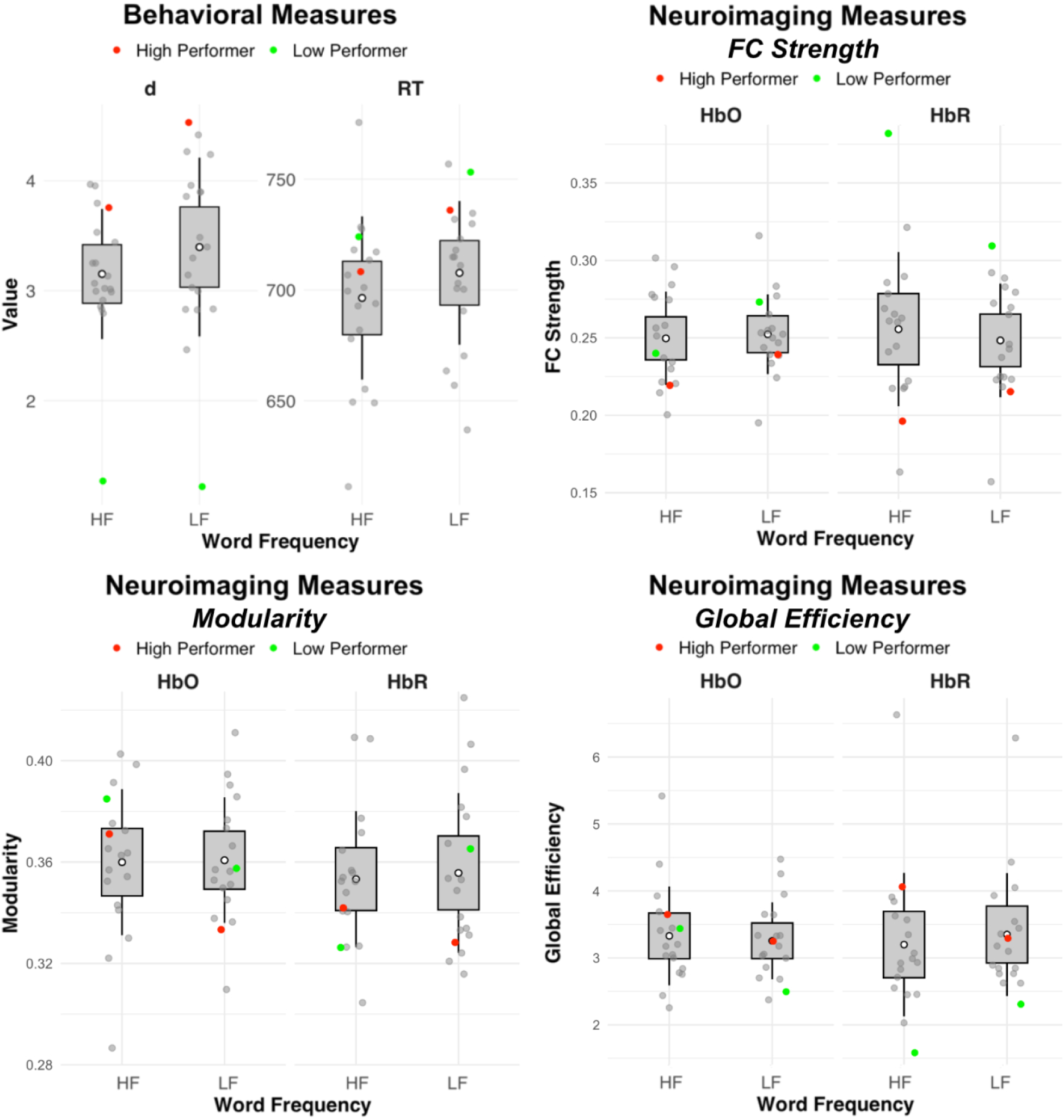
Data distribution for behavioral (*d’* and RT) and neuroimaging (FC strength, modularity and global efficiency) measures, with high and low performers highlighted in red and green, respectively. The boxplots represent the mean, 95% confidence intervals (hinges), and standard deviations (whiskers). RT = reaction time, HF = high frequency, LF = low frequency, HbR = deoxyhemoglobin, HbO = oxyhemoglobin, FC = functional connectivity.

One question is whether the lack of a main effect for word frequency across the FC measures was due to averaging over thresholds. To address this question, we asked whether there was a consistent difference between HF and LF conditions for the measures as a function of threshold (**Figure 5**). For modularity, the word frequency derived from both HbO and HbR were both significant. For modularity derived from HbO, the main effects of word frequency [Word Frequency: *β* = 0.006, SE = 0.001, *t*(3168) = 4.39, *p* < 0.001, linear [*β* = -0.447, SE = 0.013, *t*(18.74) = -33.7, *p* < .001] and quadratic threshold [*β* = 0.874, SE = 0.012, *t*(3168) = 75. 75, *p* < .001] were significant. Word Frequency * Threshold interactions were significant linear [Word Frequency * Threshold: *β* = 0.016, SE = 0.004, *t*(3168) = 4.27, *p* < .001] and the quadratic threshold [Word Frequency * Threshold: *β* = -0.077, SE = 0.016, *t*(3168) = -4.73, *p* < .001]. Similarly, word frequency was also significant for HbR derived modularity [Word Frequency: *β* = 0.005, SE = 0.001, *t*(3168) = 3.37, *p* = .001], as were all other effects [Threshold: *β* = -0.442, SE = 0.011, *t*(19.05) = -41.63, *p* < .001; Threshold^2^: *β* = 0.855, SE = 0.011, *t*(3168) = 78.29, *p* < .001]; Word frequency*Threshold: *β* = 0.008, SE = 0.004, *t*(3168) = 2.25, *p* = .025; Word frequency*Threshold^2:^ *β* = -0.033, SE = 0.015, *t*(3168) = -2.13, *p* = .033].

**Figure 5.**
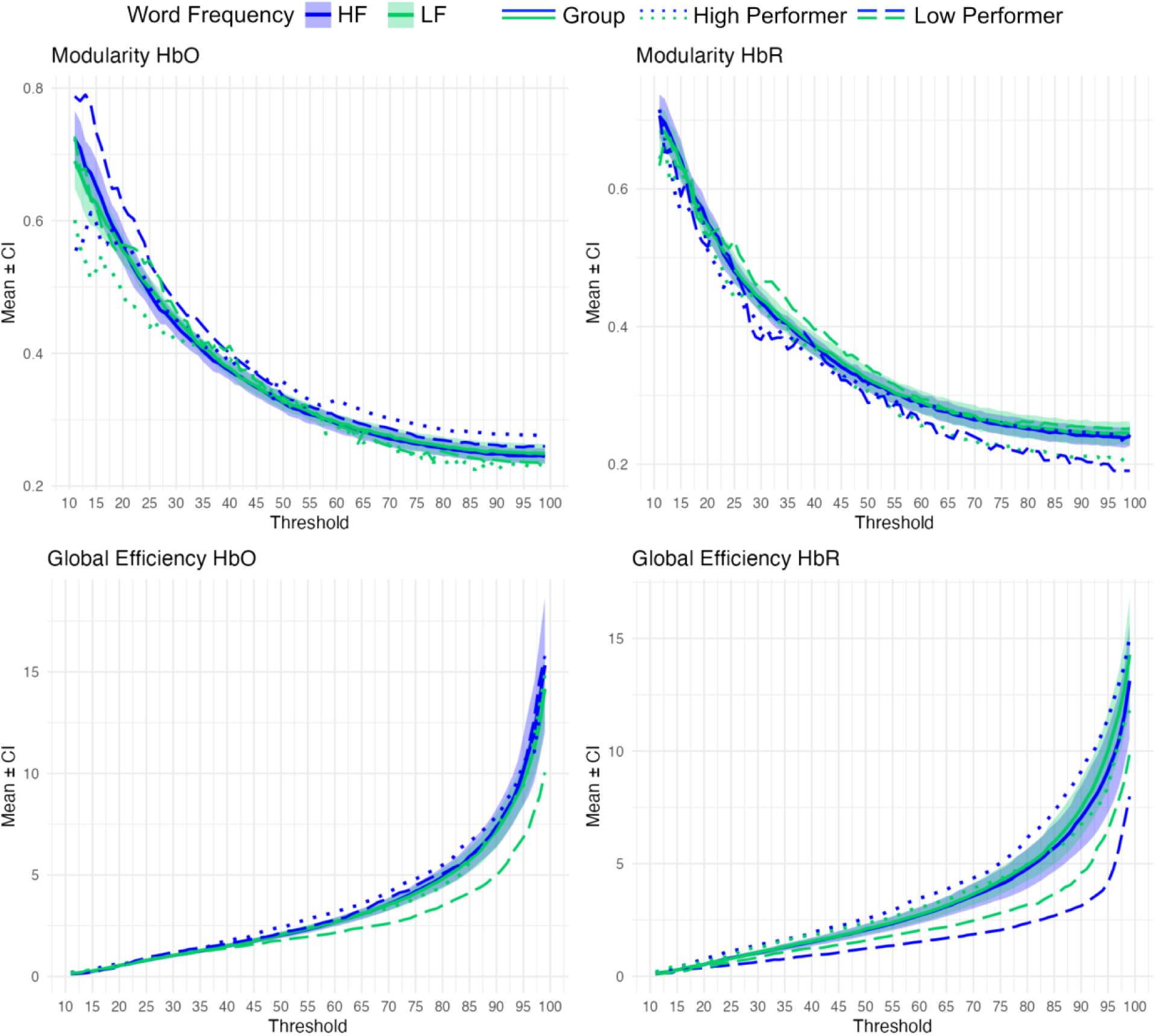
Modularity and global efficiency in the group excluding the extreme performers (solid lines representing group mean and 95% CIs) as well as the high performer (dotted lines) and the low performer (dashed lines) over thresholds (10% to 99% threshold in 1% intervals), for high and low frequency words (HF in blue, LF in green). HF=high frequency, LF=low frequency, HbR=deoxyhemoglobin, HbO=oxyhemoglobin.

Global efficiency derived from both HbO and HbR revealed Word Frequency * Threshold effects. For global efficiency derived from HbO, although there was no main effect of Word Frequency [Word Frequency: *β* = 0.06, SE = 0.066, *t*(3186) = 0.86, *p* = .39], there was a significant main effects of threshold [Threshold: *β* = 10.62, SE = 0.517, *t*(19.05) = 20.53, *p* < .001; Threshold^2^: *β* = 20.01, SE = 0.53, *t*(3186) = 37.72, *p* < .001], and a significant Word frequency*Threshold interaction [Word frequency*Threshold: *β* = -0.48, SE = 0.172, *t*(3186) = - 2.80, *p* = .005; Word frequency*Threshold^2^: *β* = -1.97, SE = 0.75, *t*(3186) = -2.63, *p* = .009]. Similarly, for global efficiency derived from HbR, although there was no main effect of word frequency [Word Frequency: *β* = -0.02, SE = 0.054, *t*(3186) = -0.38, *p* = .70], there were main effects for threshold [Threshold: *β* = 9.76, SE = 0.815, *t*(18.27) = 11.97, *p* < .001; Threshold^2^: *β* = 16.26, SE = 0.432, *t*(3186) = 37.67, *p* < .001], and the Word frequency*Threshold interactions were significant [linear: *β* = 0.76, SE = 0.140, *t*(3186) = 5.38, *p* < .001; quadratic: *β* = 2.6, SE = 0.61, *t*(3186) = 4.25, *p* < .001]. As can be seen in Figure 5, both modularity and global efficiency derived from both HbO and HbR indicates that LF words are associated with greater modularity and global efficiency as compared to HF words across the thresholds.

### Relationship Between Behavioral and Functional Connectivity Measures

To understand the relationship between FC measures and cognitive function, we also examined the correlation between performance accuracy, and the FC network measures (**Figure 6**). For the HbR derived signal, we observed that global efficiency was positively correlated with *d’* for the high frequency words [*R* = 0.46, *p* = .027] with greater *d’* for being linked to higher GE. GE was not correlated with *d’* for low frequency words [*R* = 0.09, *p* = 0.72]. There was also a negative relationship between RT and global efficiency for HF words [*R* = -0.43, *p* = .039], but not LF words [*R* = 0, *p* = 0.99]. There were no significant effects related to for the HbO derived signal. There were also no significant or marginal relationships between the modularity, FC strength and task performance. Overall, the results suggest that a highly efficient prefrontal network is beneficial for task performance, but the effect is specific to HF words.

**Figure 6.**
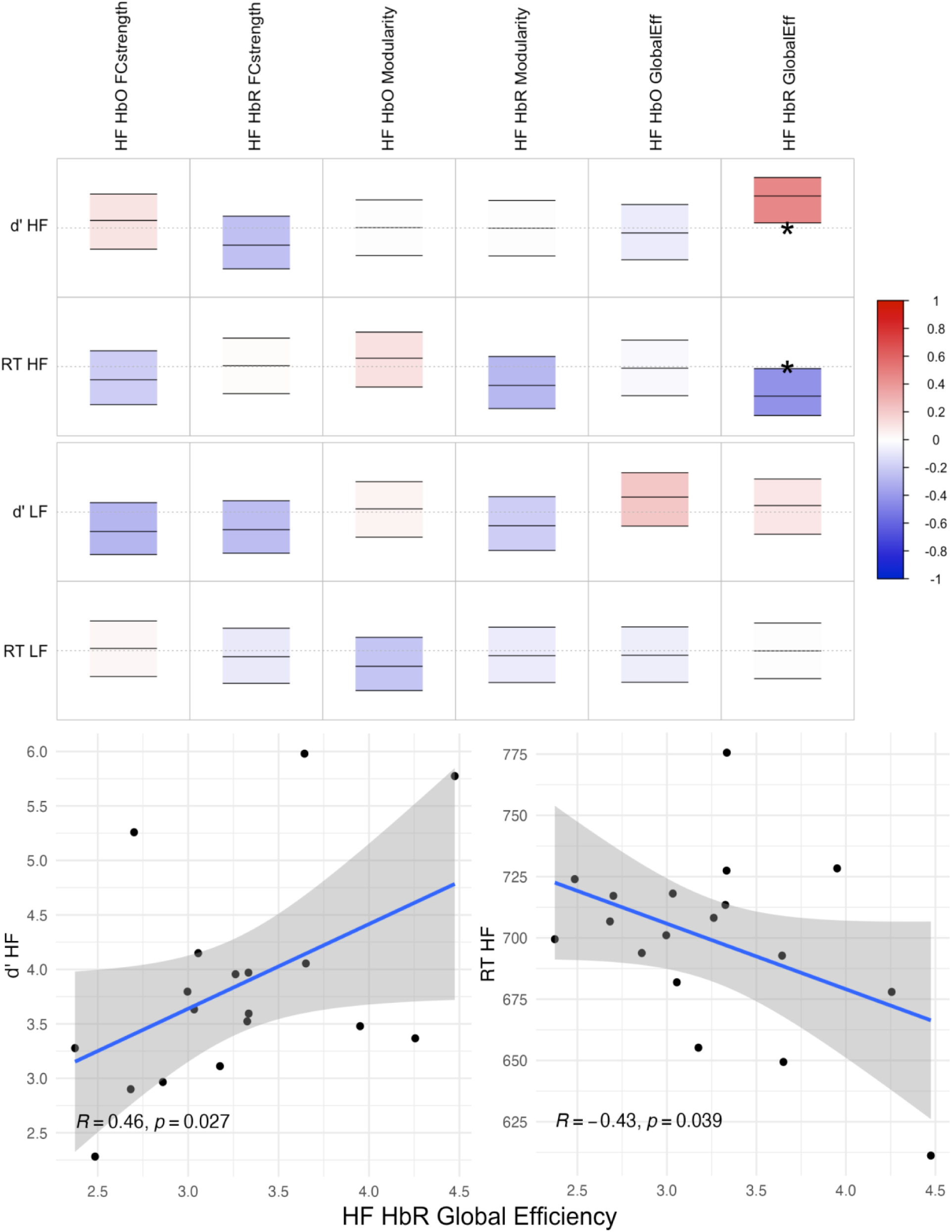
Brain-behavior relationships. Correlations between task performance (*d’*, RT) and functional connectivity measures of strength, modularity, and global efficiency, for the high word frequency (top) and low word frequency (bottom) tasks. Each cell of the correlation matrix contains the correlation coefficient (middle line) bound by the upper and lower 95% confidence intervals, with the color of each rectangle indicating the direction of the relationship (* *p* < .05). The relationship between *d’* and global efficiency (HbR signal) during the HF task was significant before but not after multiple comparison correction (see scatterplot), so that higher global efficiency was associated with better task performance.

In addition to accuracy and RTs changing significantly over the course of the blocks, as can be seen in **Figure 3**, there was a notable range in the performance across the individual participants. For the participant having the highest accuracy (red), RTs were also well above the normal range (**Figure 3**) for both the HF and LF conditions. In constrast, RTs for the participant have the lowest performance accuracy (green) fell within the normal range or both the HF and LF conditions. Since our global efficiency was significantly correlated with accuracy and RTs, we next asked if our graph theory approach was also sensitive to individual differences. We calculated individual z-scores for the behavioral and functional connectivity measures (See **Table 2**) for the highest performing (High-P) and the lowest performing participant (Low-P) to examine whether their connectivity profiles resemble each other’s as well as those of the rest of the group.

**Table 2.** Means and standard deviations for behavioral and brain measures & *z*-scores for Highest and (High-P) and Lowest Performing (Low-P) Participants.

| Behavioral Measures |  |  |  |  |
| --- | --- | --- | --- | --- |
| ***d'/RT*** | ***d' HF*** | ***d' LF*** | ***RT HF*** | ***RT LF*** |
| **Mean** | 3.150 | 3.400 | 696.36 | 707.69 |
| **SD** | 0.590 | 0.810 | 36.87 | 32.42 |
| **High-P z-score** | **1.155** | **1.503** | 0.330 | 0.916 |
| **Low-P z-score** | **-4.961** | **-3.708** | 0.781 | **1.532** |
| Graph Theory Measures |  |  |  |  |
| ***FC strength*** | ***HF HbO*** | ***LF HbO*** | ***HF HbR*** | ***LF HbR*** |
| **Mean** | 0.250 | 0.252 | 0.256 | 0.248 |
| **SD** | 0.030 | 0.026 | 0.050 | 0.037 |
| **High-P z-score** | **-1.067** | -0.523 | **-1.257** | -0.878 |
| **Low-P z-score** | -0.326 | 0.838 | **3.578** | **1.718** |
| ***Modularity*** | **HF HbO** | **LF HbO** | **HF HbR** | **LF HbR** |
| **M** | 0.360 | 0.361 | 0.353 | 0.356 |
| **SD** | 0.029 | 0.025 | 0.027 | 0.032 |
| **High-P z-score** | 0.496 | -0.978 | -0.417 | -0.886 |
| **Low-P z-score** | 0.991 | 0.009 | **-1.052** | 0.333 |
| ***Global Efficiency*** | **HF HbO** | **LF HbO** | **HF HbR** | **LF HbR** |
| **M** | 3.329 | 3.255 | 3.199 | 3.349 |
| **SD** | 0.738 | 0.575 | 1.071 | 0.919 |
| **High-P z-score** | 0.433 | 0.012 | 0.810 | -0.013 |
| **Low-P z-score** | 0.191 | **-1.460** | **-1.558** | **-1.257** |

Note. RT = reaction time, HF = high frequency + phonotactic probability, LF = low frequency + phonotactic probability, HbO = oxyhemoglobin signal, HbR = deoxyhemoglobin signal.

The highest performing participant showed low FC strength for the HF task condition in both signals, whereas the lowest performing participant showed high FC strength for HF and LF words in the HbR signal, relative to the rest of the cohort. With respect to modularity, the highest performing participant did not show divergence form the rest of the cohort in terms of modularity whereas the lowest performing participant showed low modularity during the HF task for the HbR signal. Finally, for global efficiency, again, the high performer did not differ from the rest of the cohort. Overall, the high performer’s prefrontal network did not show a non-typical organization but supported a lower than FC strength. In contrast, the low performer showed low global efficiency during the HF task, observed in the HbR signal and the LF task for both types of signals. Further, the low performer’s network had low global efficiency alongside high FC strength. Relating the *z*-scores to the correlations reported in the previous section, we observed that in the cohort low global efficiency was related to low *d’* and marginally high RT for the HF task and HbR signal, consistent with the trend observed in the entire sample.

## Discussion

This is the first study to apply a graph theoretical approach to the investigation of FC of brain networks using HbO and HbR fNIRS signals in typical language participants. We characterized the connectivity profiles using FC and graph theory measures and found them to be sensitive to individual differences in verbal working memory. Participants showed a consistent difference in modularity between HF and LF words, presenting a consistently higher modularity for low frequency words as compared to high frequency words.

### Efficiency of language processing is affected by word frequency

In the study, we compared the behavioral performance and brain measures of FC strength, modularity and global efficiency for HF and LF words in a typical language sample. While there were no significant main effects of word frequency in the behavioral analysis, we observed that the participants became more accurate but slower across subsequent task blocks for the LF blocks as compared to the HF blocks, replicating our findings the from Berglund-Barraza, et al., 2019, 2020 for RT but with smaller effect sized for accuracy due in part to the non-overlapping participant sample in the current study.

We also observed greater modularity (e.g., lower network integration) for the LF words as compared to the HF words; however, the effect was not consistent over the threshold range due to the presence of the significant interactions. Nevertheless, this might suggest that LF words were easier to process as compared to HF, mirroring the accuracy findings from *d*’. Optimal modularity depends on the cognitive context, where dominance of local, segregated processing is favored at rest and during easy cognitive tasks, while higher integration due to need for inter-module information exchange is required as cognitive task demands grow. For example, in an fMRI study of typical young adults (Cohen & D’Esposito, 2016) showed lower modularity (or greater network integration) during the n-back task compared to a motor sequence tapping task and resting-state. Such greater network integration was correlated with high accuracy on the *n*-back task.

Together, the results from this study show that words that have both high in overall frequency of occurrence in the lexicon, and which are also composed of phonemes whose order of occurrence in the English language are also highly frequency posed a greater challenge to processing, updating and actively maintaining spoken on the part of the participants as compared to word having low overall frequency and low phonotactic probability. This is consistent with the prediction of spoken word recognition models that argue that *lateral inhibition* connections enable words that have strong activations to directly suppress less-active competitors and that the degree of lateral inhibition is what makes the recognition of a give word more or less efficient (Gur & McMurray, 2025). Although these models focus on the processing of spoken words in isolation, taken together with the finding from this study suggest that when scaling up to spoken sentence comprehension where the individual must process, update and actively maintain multiple words in memory over the course of the sentence, the individual features of words carry unique processing demands that be accumulative over the course of the sentence.

Of note, this word frequency effect on modularity was not seen when averaging the measure across all thresholds, possibly suggesting that networks that are too sparse or too dense might not be able to capture differences in modularity between task conditions. However, it is not clear what the appropriate threshold values are, necessitating testing across a range of threshold values.

### Efficiency of language processing differs between typical individuals

This study also characterized the FC patterns of participants with low and high task performance relative to the rest of the sample. We observed that the high performing individual tended to show low FC strength, while the low performer showed high FC strength. This measure might be interpreted as total network integration. High values of FC strength might therefore represent indiscriminate and maladaptive attempts at establishing a connectivity pattern which should maximize efficiency of information transfer while minimizing energetic cost. While this effect was observed in an otherwise typical individual, inefficient FC patterns have been previously described in multiple neurological disorders, including those on the side of neurodegeneration (Chen et al., 2021; Yao et al., 2010) as well as proposed to arise with atypical brain network development, such as schizophrenia (Lynall et al., 2010) or autism spectrum disorder (Rudie et al., 2012).

Furthermore, we observed that the low performing individual showed low modularity, and even more so, low global efficiency. *Modularity* captures the strength of segregation between non-overlapping functional modules. This modular organization is driven by economical constraints driving minimization of connection cost (Achard & Bullmore, 2007; Bullmore & Sporns, 2012) and ensures efficient information transfer in the entire network through a balance of specialized local processing and longer connections facilitating global integration (Rubinov & Sporns, 2011; Bertolero et al., 2015). On the other hand, global efficiency captures the capacity of a network to integrate information by quantifying how efficiently information can be exchanged between all pairs of nodes through the shortest communication paths. Global efficiency is considered a key measure of functional integration using long-range connections, reflecting the network’s ability to support coordinated information processing across the whole network (Bullmore & Sporns, 2009).

Our findings support those of Cohen & D’Esposito (2016) who showed lower modularity (or high integration) was correlated with higher accuracy on the *n*-back task. The cited study assessed the modularity of the entire brain composed of multiple distinct networks which might need to integrate to tackle high cognitive task demands.

### Methodological considerations of fNIRS

These findings suggest that fNIRS provides the ability to register the optical signal of fNIRS to neuroanatomical data, making capturing the spatial temporal dynamics of spoken language and cognitive processing both feasible and cost-effective (Schroeder et al., 2023; Telkemeyer et al., 2009). As a result, fNIRS is gaining significant traction as a versatile optical neuroimaging tool to assess auditory paradigms and language function in both typical hearing listeners, those with developmental language disorders, aging populations, and those fitted with hearing aids and cochlear implants (Sherafati et al., 2022). It is particularly suitable for investigation of auditory paradigms due to its silent operation, high degree of compatibility with hearing device use, higher spatial resolution than EEG, better temporal resolution than fMRI, low sensitivity to motion artifacts, ease of use with children, and individuals with autism spectrum disorder. Both oxygenated (HbO) and deoxygenated (HbR) hemoglobin exhibit strong but distinct oxygen-dependent optical absorption, so that the concentration of HbO and HbR in the brain can be measured using near-infrared light (Eggebrecht et al., 2012, 2014). Changes in local blood oxygenation including HbO and HbR serve as proxy for experimental stimulus-dependent neural activity (Yücel et al., 2021).

### Limitations

While fNIRS technology offers many advantages, it does have its limitations, especially when using sparse probe arrays. Despite higher temporal resolution, its spatial resolution and signal-to-noise ratio (SNR) are relatively low compared to fMRI. Furthermore, its coverage is limited to ∼2-2.5 cm of the cortical surface, with no access to deep sulci and the subcortex. The functional connectivity measured from a sparse probe array were constrained by its limited spatial coverage, resulting in the observation of the incomplete large-scale networks, which are typically estimated across the entire brain when using fMRI. The network nodes were dictated by the layout of the probe and might not represent the typical hub regions which would be chosen using a whole-brain method. Additionally, despite preprocessing there might be residual physiological contamination and motion artifacts. These limitations can be ameliorated by using high-density diffuse optical tomography (HD-DOT) systems, an optical method with superior spatial resolution, allowing for image reconstruction with depth information, and therefore better coverage of the distributed cortical networks. Therefore, future studies should aim to use HD-DOT for more reliable functional connectivity estimates that are challenging to interpret using sparse array fNIRS.

### Conclusions

In this study, we showed that the prefrontal brain networks of participants adapt to differing demands of processing, updating and actively maintaining spoken words that differ in spoken word frequency and phonotactic probability, as indicated by the consistent differences in modularity between high and low frequency words. Crucially, we have shown that even in typical development and young adulthood, individual differences in cognitive performance are related to network properties capturing processing efficiency. This is also the first study applying the graph theoretical approach to the investigation of FC of brain networks using fNIRS to characterize the multi-domain and language network engagement. The results are robust across many thresholds, providing preliminary methodological guidelines for future research using this method. Further investigation of graph measures might reveal qualitative differences between participants as well as among different clinical populations, including DLD, ASD, and others.

## DATA AVAILABILITY STATEMENT

The datasets generated for this study are available on request to the corresponding author.

## ETHICS STATEMENT

All participants completed the written informed consent protocols in accordance with the Declaration of Helsinki as well as the guidelines of the University of Texas at Dallas Institutional Review Board (IRB), which approved the protocol.

## AUTHOR CONTRIBUTIONS

JE designed the experiment, developed the stimuli. Along with doctoral students in the lab, JE also collected, processed and analyzed the data. PS conducted the functional connectivity and graph theory analysis. AE provided expertise in the analysis of the fNIRS data. JE and PS jointly wrote and edited the manuscript. AE provided feedback on the writing of the manuscript.

## FUNDING

This study was funded by the University of Texas at Dallas research funds to JE and the National Institute on Deafness and Other Communication Disorders R01-DC005650 and K18-DC021149 to JE.

## ACKNOWLEDGMENTS

The authors gratefully thank Dr. Berglund-Barraza and the young adults who participated in the research.

## Notes

### Competing Interest Statement

The authors have declared no competing interest.

### Summary of Updates

The manuscript has been revised following the co-authors' comments. The data was re-analyzed using a more fitting filter. We removed descriptive and non-significant results that detracted from the main narrative of the paper. The Figures were edited for clarity. The introduction has been rendered more concise.

